# Wing Geometry and Genetic Analyses Reveal Contrasting Spatial Structures between Male and Female *Aedes aegypti* Populations in Metropolitan Manila, Philippines

**DOI:** 10.1101/2020.09.16.299487

**Authors:** Thaddeus M. Carvajal, Divina M. Amalin, Kozo Watanabe

## Abstract

**Background:** Many important arboviral diseases (e.g. dengue, chikungunya) are transmitted by the bite of a female mosquito vector, *Aedes aegypti*. Hence, the population genetic structure of the mosquito has been studied in order to understand its role as an efficient vector. Several studies utilized an integrative approach; to combine genetic and phenotypic data to determine the population structure of *Ae. aegypti* but these studies have only focused on female populations. To address this particular gap, our study compared the population variability and structuring between male and female *Ae. aegypti* populations using phenotypic (wing geometry) and genetic (microsatellites) data from a highly-urbanized and dengue-endemic region of the Philippines, Metropolitan Manila.

**Methods:** Five mosquito populations comprised of female (n = 137) and male (n = 49) adult *Ae. aegypti* mosquitoes were used in this study. All mosquito individuals underwent geometric morphometric (26 landmarks), and genetic (11 microsatellite loci) analyses.

**Results:** Results revealed that *F_ST_* estimates (genetic) were 0.055 and 0.009 while *Q_ST_* estimates (phenotypic) were 0.318 and 0.309 in in male and female populations, respectively. Wing shape variation plots showed that male populations were distinctly separated from each other while female populations overlapped. Similarly, discriminant analysis of principal components using genetic data revealed that male populations were also distinctly separated from each other while female populations showed near-overlapping populations. Genetic and phenetic dendrograms showed the formation of two groups in male populations but no groups in female populations. Further analysis indicated a significant correlation *(r* = 0.68, *p* = 0.02) between the genetic and phenetic distances of male populations. Bayesian analysis using genetic data also detected multiple clusters in male (K = 3) and female (K = 2) populations, while no clusters were detected using the phenotypic data from both sexes.

**Conclusions:** Our results revealed contrasting phenotypic and genetic patterns between male and female *Ae. aegypti*, indicating that male populations were more spatially structured than female populations. Although genetic markers demonstrated higher sensitivity in detecting population structures than phenotypic markers, correlating patterns of population structure were still observed between the two markers.

## Background

*Aedes aegypti* is the primary mosquito vector for several important mosquito-borne diseases such as dengue, chikungunya and Zika. Over the past decade, many scientists have focused on studying the population genetic structuring of this species within urban areas^1-6^. Genetic markers such as microsatellite loci and single nucleotide polymorphisms (SNPs) have been largely used to investigate the population structure of *Ae. aegypti* on macro- and micro-geographic scales which revealed high genetic diversity and distinct genetic clustering in different regions and countries^7-9^, cities and villages^1,2,6^.

Wing geometry is a phenotypic marker that can be used as an alternative approach to describe the population variability and structure of *Ae. aegypti* since these are evolutionarily informative and heritable^10,11^. Previous studies on *Ae. aegypti* have demonstrated that wing shape can be an indicator of population genetic structure in fine-scale areas^12,13^. It can also detect subtle variation within a single mosquito population either over time (e.g. temporal variation)^4^ or along environmental gradients (e.g. altitude, levels of urbanization) ^14,15^. For this reason, estimating the population genetic structure using wing geometry has been supported by many studies because of its low-cost alternative^10^.

Many independent studies have focused on either genetic or phenotypic markers, but some several studies have also integrated both markers^16-19^, especially in *Ae. aegypti*^3,4^ Patterns of wing shape variation and genetic diversity in *Ae. vexans* indicated distinct spatial structural differences from northern and central European countries with considerable gene flow on a regional scale^19^. Parallel temporal changes in allelic frequencies and wing shape were also observed in *Ae. aegypti* from Brazil, suggesting that these changes could be driven by genetic drift and divergent selection^4^, however, the results generated by genetic and phenotypic markers have often contradicted each other. For example, estimates of the morphological diversity index (*Q*_ST_) have often been larger than the genetic diversity index (*F*_ST_) from mosquito populations at the micro-^3,4^ and macro-geographical^16^ scale in Brazil. Wing shape was also unable to show clear patterns of population differentiation compared to genetic markers at these scales^3,18^. The slow evolutionary rate of change for wing shape which has resulted in morphological uniformity or homogeneity could be due to high gene flow or continuous migrations of this mosquito vector that counteract local genetic drift^10,20^. More importantly, since wings of *Ae. aegypti* are important organs for flight and sexual signaling, selective pressure may have stabilized them over time^21^.

The majority of these integrative (wing geometry and genetic) analysis investigations only focused on female *Ae. aegypti* populations. This may be due to the importance of female mosquitoes in transmitting arboviruses, but research focusing on male *Ae. aegypti* mosquitoes is becoming equally important due to its notable role in vector control strategies, particularly in rear-release methods such as sterile insect technique (SIT), insect incompatibility technique (IIT) and genetically modified mosquitoes (GMM)^22-24^. Notable biological differences in male and female *Ae. aegypti* mosquitoes could affect genetic and phenotypic variability as well as the spatial structuring of each sex. For instance, smaller-sized male mosquitoes may have shorter dispersal distance, thereby producing heterogeneous male populations even on the micro-geographic scale. This was exemplified in our previous wing geometry study^12^ which revealed the fine-scale population structure in male mosquitoes with short dispersal distances (up to 22 km).

The aim of this study was to describe and compare the population variability and structure between male and female *Ae. aegypti* mosquitoes using wing geometry and genetic markers. Adult *Ae. aegypti* mosquitoes were collected from within a highly-urbanized and dengue-endemic region of the Philippines, Metropolitan Manila. Eleven microsatellite loci and 26 identified morphometric landmarks were used to compare both sexes. Unlike previous studies that used different mosquito individuals for each approach^3, 16-18^, we utilized the same mosquito for genetic and morphometric analysis in order to reflect and compare the population variability and structure for both markers.

## Methods

### Study Area and Mosquito Sampling

Collection of *Ae. aegypti* adult mosquitoes was conducted within Metropolitan Manila, Philippines (Figure 1) from May 2014 to January 2015. Households were selected based on voluntary consent to collecting adult mosquitoes on their premises. The collection procedure, sorting, preservation and identification of adult mosquitoes was based on Carvajal et al.^6^ while sex determination of each adult was performed using the morphological pictorial keys from Rueda et al.^25^.

**Figure 1.**
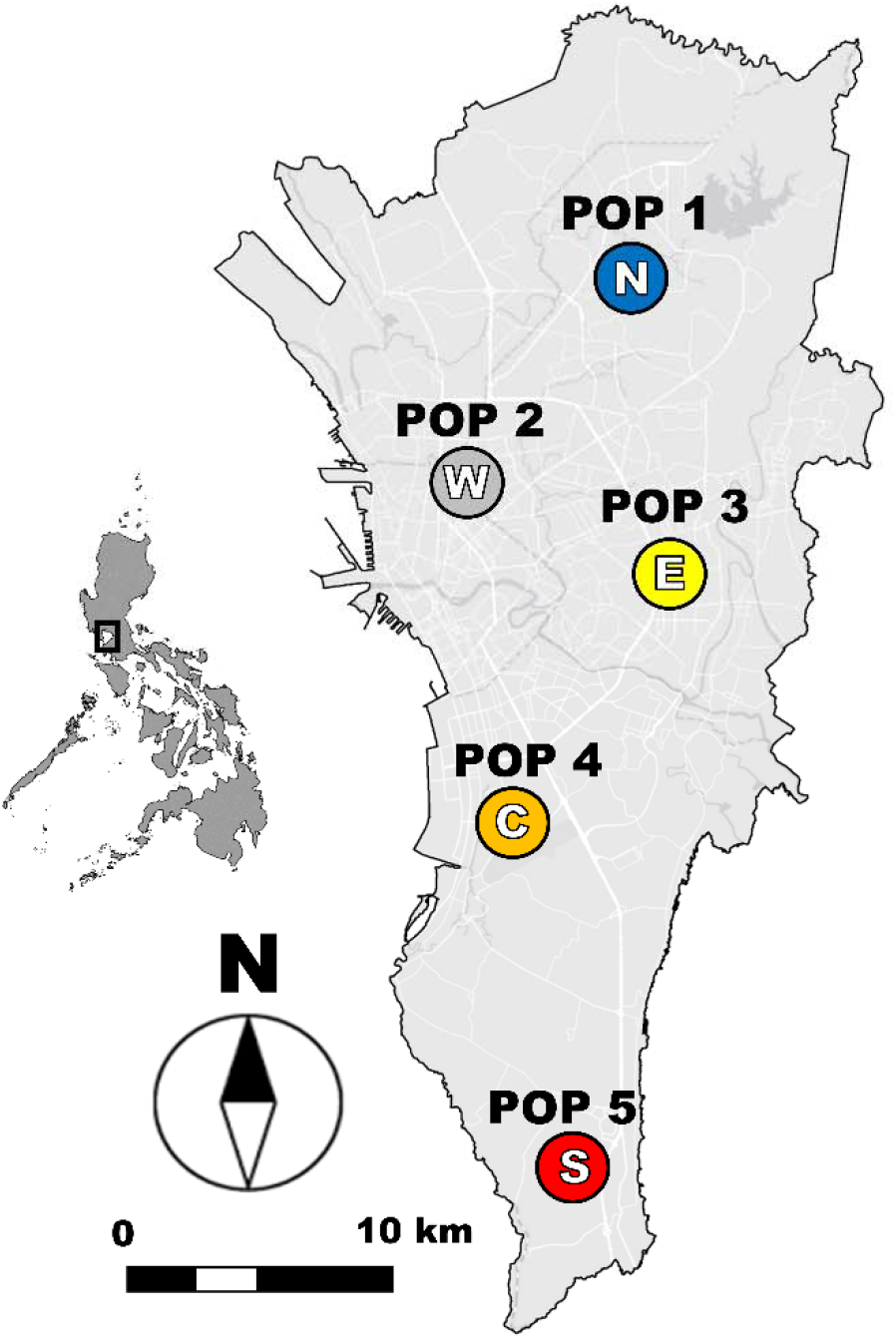
Geographic midpoints of *Ae. aegypti* mosquito populations in Metropolitan Manila. Colored circles represents each mosquito population: POP 1 (N = North), POP 2 (W = west), POP 3 (E = east), POP 1 (C = central) and POP 1 (S = south). This map was prepared using the ArcGIS version 10.2.2 from LandsatLook Viewer (http://landsatlook.usgs.gov/)

In this study, five mosquito populations (POP) represented the entirety of Metropolitan Manila: POP 1 (north), POP 2 (west), POP 3 (eastern), POP 4 (central) and POP 5 (south) (Table S1). A total of 66 households participated in the study and the number of households ranged from 9 – 16 for female mosquitoes and 3 – 5 for male mosquitoes, respectively (Table S1). Since households were widely dispersed, a geographical midpoint^26^ was calculated to assign a single georeferenced location for each mosquito population. A total of 186 individuals consisting of 137 females and 49 males were used for subsequent phenotypic and genetic analyses (Table S1).

### Geometric Morphometric Analysis

Both wings were detached from the thorax of each adult mosquito by carefully cutting at the base of the attachment with a sterilized blade and forceps. Afterwards, the remaining portions of each mosquito were placed in a microcentrifuge with 99% ethanol for genetic processing. Wing scales were removed using the methods published Carvajal et al.^12^ and were then mounted on a glass slide and cover slip with Aquatex^®^ (Merck, Darmstadt, Germany). Wing images were captured with 3.5x magnification using Nikon SMZ800 dissecting microscope. The number of landmarks used by Carvajal et al.^12^ was maintained when digitizing each mosquito wing using TpsDig V2.10^27^.

Wing size was assessed using the centroid size, an isometric estimator, from the landmark coordinates computed from MorhoJ^28^. A comparative analysis of the centroid sizes was performed for populations of both sexes using parametric Analysis of Variance (ANOVA) and between males and females using an independent *t*-test in P.A.S.T. software^29^. A test of allometry was initially performed to determine the effect of wing size on wing shape through multiple regressions of the Procrustes coordinates (shape) and log-transformed centroid sizes with 10,000 permutations. The Generalised Least-Squares Procrustes superimposition algorithm^30^ was used to produce the wing shape variables (partial wraps). Afterwards, discriminant analysis was performed on test for dissimilarities between mosquito populations. This analysis generated canonical variates (CV) which can be represented as a scatterplot (morphospace), and also calculated pairwise Mahalanobis distances among mosquito populations with 10,000 permutations. All these analyses were done using the MorhoJ software ^28^. To statistically validate the comparisons, the significance of the metric disparity (MD) of partial wraps between mosquito populations and the quantitative estimator of intra/inter population differentiation (*Q*_ST_) were tested by permutation tests (2,000 iterations each) using COV software^31^. A phenotypic dendrogram was constructed based on the Mahalanobis distance using the unweighted pair group with arithmetic mean (UPGMA) in the *fastcluster* package^32^ in R version 3.5^33^.

### Genetic Analysis

The microsatellite data of selected mosquito samples were obtained from a previous population genetics study^6^ deposited at vectorbase.org^34^. DNA extraction, multiplex PCR methods, fragment analysis and genotyping of 11 selected microsatellites were based on the protocol provided Carvajal et al.^6^. Microsatellite data error and presence of null alleles were examined using MICROCHECKER^35^. Hardy-Weinberg equilibrium (HWE) tests and estimations of the linkage disequilibrium (LD) among all pairs of loci were conducted using GENEPOP v4.2.1^36,37^. Significance levels for multiple testing were corrected using the Bonferroni procedure. Observed heterozygosity (*Ho*) expected heterozygosity (*He*), and inbreeding coefficients (*F*_IS_) for each mosquito population and analysis of molecular variance (AMOVA) were calculated using the Genetic Analysis in Excel (GenAlEx) version 6.3^38^. Pairwise *F*_ST_ values were calculated using Arlequin v3.5.1.3^39^ with 10,000 permutations to determine the magnitude of the genetic differentiation among mosquito populations. The Cavalli Sforza and Edwards distance was calculated using the FreeNA program^40^ and was used to create a genetic dendrogram based on UPGMA in the *fastcluster* package^32^ of R version 3.5^33^. To visualize patterns of genetic differentiation among mosquito populations, discriminant analysis of the principal components (DAPC) were implemented in the *adegenet*^41^ package in R version 3.5^33^ was used.

### Detecting Population Structure

To detect the population structure for wing geometry and genetic data for male and female populations, we used the Bayesian clustering algorithm from GENELAND 4.0.3^42^ in R version 3.5^33^ which accounts for the spatial location of mosquito samples to estimate the optimal number of clusters (K). This software was used because it can handle both phenotypic and genetic data in inferring the number of clusters (K) and allows for the comparative analysis of genetic and phenotypic data. The datasets cover genetic data in the form of individual genotype alleles while the morphometric data consisted of individual CV scores and georeferenced coordinates per mosquito individual. We implemented 10 replicates per 1 million iterations (thinning = 200) with the number of possible clusters *K* ranging from 2 – 5. The best probable number of clusters was determined by the highest posterior density from all preliminary runs. A final run was processed on a landscape of 200 × 200 cells with a burn-in of 2000 iterations^19,42,43^.

### Comparison and Correlation between Genetic and Phenotypic Markers

To determine if the *Q_ST_* estimate was not equal to *F_ST_* estimates, we performed a one-sample Wilcoxon test for the male and female populations. Spearman’s rank correlation coefficient between phenotypic (Mahalanobis) and genetic (Cavalli Sforza and Edwards) distances among mosquito populations was also tested. These analyses were performed using the *stats* package of R version 3.5^33^.

## Results

### Wing Size and Shape Analyses

Table S2 shows the measured centroid size of the wing for each mosquito population in male and female *Ae. aegypti*. The centroid sizes among these populations were significantly different in males (F = 3.20, p = 0.0216) than in females (F = 0.66, p = 0.6235). The average centroid sizes of the wing between female and male mosquitoes were significantly different (t = −5.8715, p < 0.001), indicating that female mosquitoes have larger-sized wings compared to male mosquitoes. Moreover, the proportion of shape variance that was explained by size (allometry) was found to be 5.43% and 1.48% in the male and female populations respectively. This demonstrates a relatively weak effect of size on shape variance.

The overall *Q*_ST_ was estimated to be 0.318 and 0.309 in the male and female populations, respectively. Metric disparity (MD) scores showed only few significantly differentiated pairwise comparisons in the male (n = 3 out of 10) and female populations (n = 4 out of 10), respectively (Table S3). Shape variation (CV) plots show that female populations overlapped with each other (Figure 2a) while male populations were distinctly separate (Figure 2b). In addition, all pairwise comparisons (Mahalanobis distance) in male and female populations were significantly differentiated.

**Figure 2.**
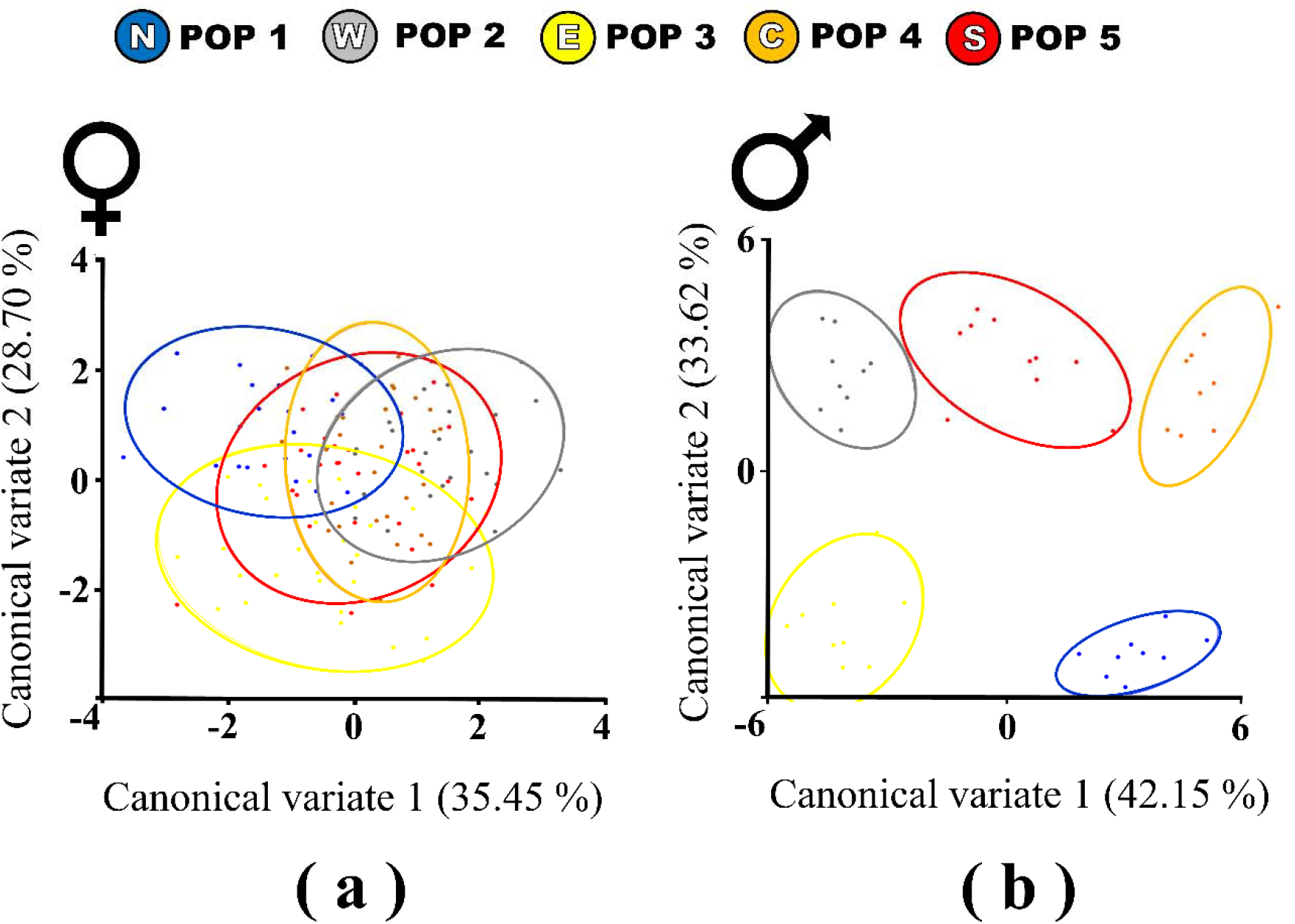
Scatterplots of the shape variation (canonical variates) in (a) female and (b) male *Ae. aegypti* populations. Colored circles represent each mosquito population (POP 1-5) from the north (N), west (W), east (E), central (C) and south (S).

Figure 4a and 4c show the phenotypic dendrograms of male and female populations based on the Mahalanobis distance. The dendrograms revealed no distinct phenotypic groups in female populations but two phenotypic groups were formed in male populations. The first group in the male phenetic dendrogram consisted of mosquito populations from the west (POP2), east (POP3) and south (POP5) while the second group consisted of north (POP 1) and central (POP 4) populations. Bayesian analysis in GENELAND detected no phenotypically structured clusters (*K* = 1) in both male and female individuals (Figure S1).

### Genetic Analysis

We observed a total of 83 and 86 alleles across 11 microsatellite loci in the male and female *Ae. aegypti* populations, respectively. The average number of alleles per loci was 7.54 in the male populations and 7.82 in the female populations (Table S4). The percentage of loci that deviated significantly from the Hardy-Weinberg Equilibrium expectations after sequential Bonferroni correction were 18.18% (male) and 38.18% (female) (Table S5). Null alleles were present on one locus (AC4) in the male mosquito populations and on two loci (AG7,B07) in the female mosquito populations (Table S6). The LD tests showed that 1.82% (male) and 6.91% (female) loci pairs had significant LD after Bonferroni corrections with no repetitions of pairs of loci combinations across mosquito populations (Table S7).

Table S8 shows the summary of the genetic diversity of male and female mosquito populations. Global *F*_ST_ estimates were 0.055 in male and 0.009 in female populations. Pairwise *F*_ST_ values in male mosquito populations ranged from 0.000 to 0.062 while female populations showed a range of 0.000 to 0.020 (Table S9). Furthermore, the results from AMOVA showed significant genetic differentiation among populations (*F*_ST_) in males (*p* < 0.05) but not in females (Table S10). Similarly, the DAPC plots (Figure 3) revealed that the male populations were distinctly separated from each other while, the female populations showed near-overlapping populations.

**Figure 3.**
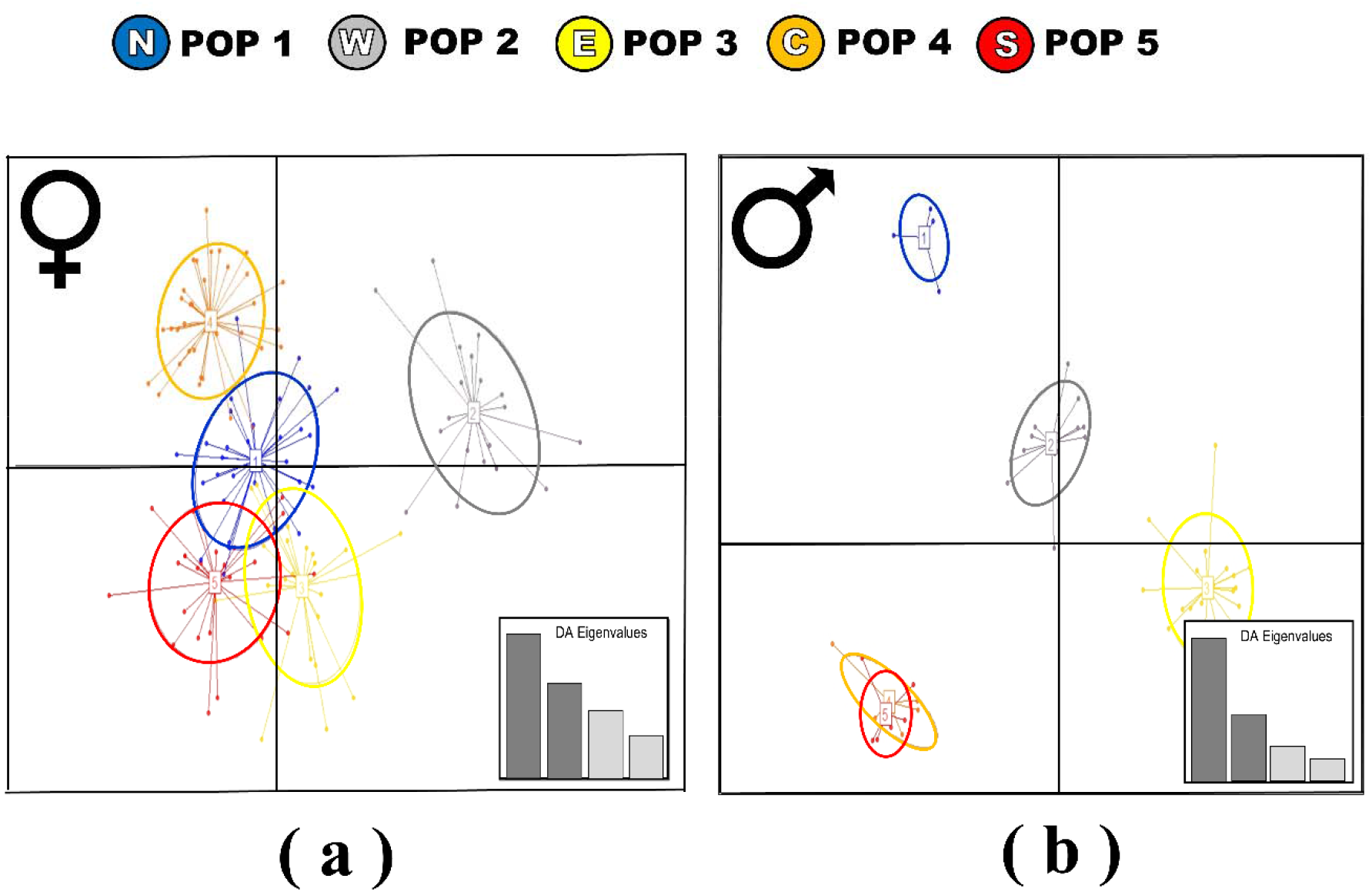
Discriminant analysis scatterplots using principal components Analysis (DAPC) in (a) male and (b) female *Ae. aegypti* populations. Colored circles represent each mosquito population (POP 1-5) from the north (N), west (W), east (E), central (C) and south (S).

**Figure 4.**
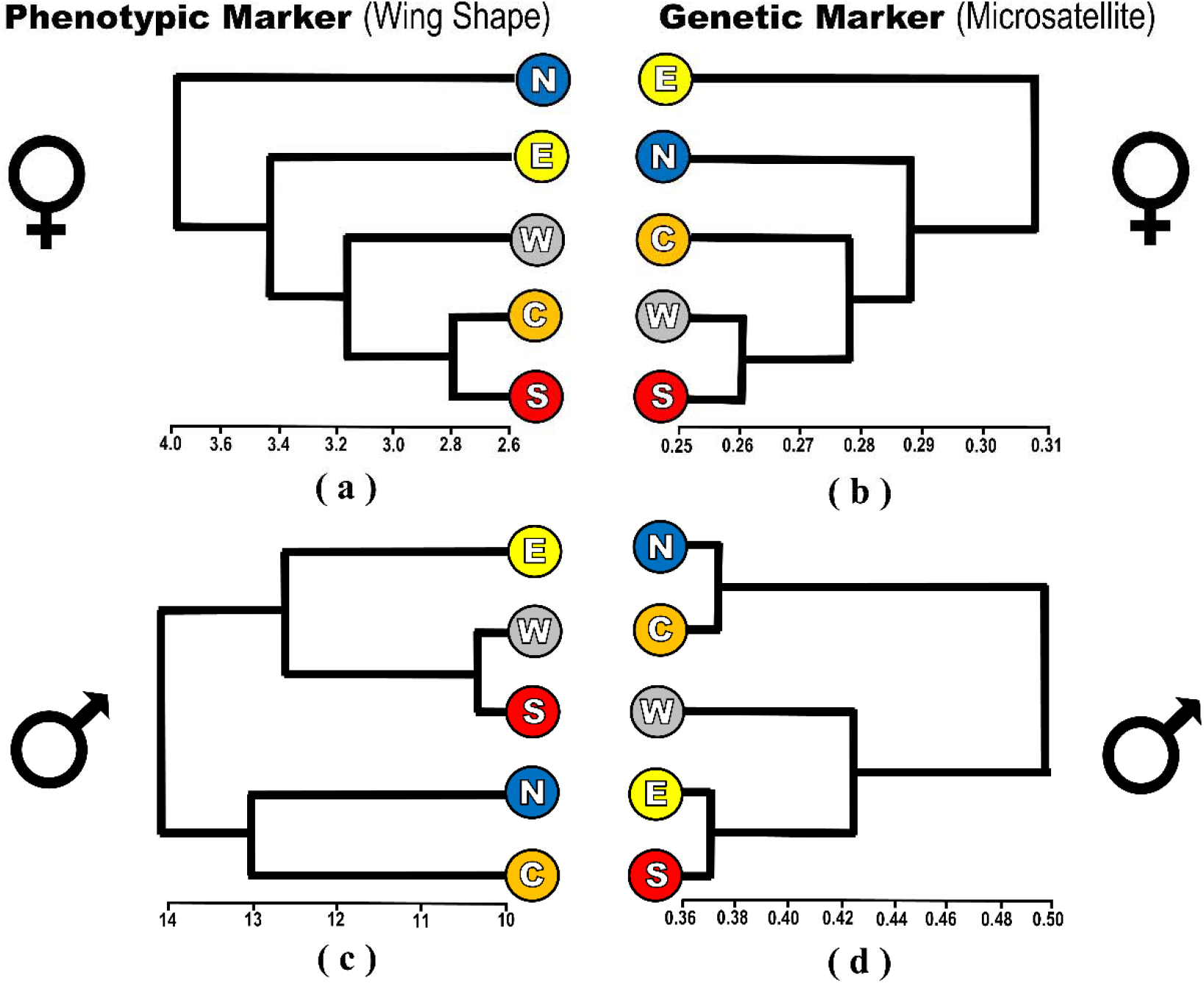
Dendrograms of *Ae. aegypti* mosquito populations; (a) phenotypic and (b) genetic dendrograms in females, (c) phenotypic and (d) genetic dendrograms in males. Mosquito populations are labeled as N (POP 1, north), W (POP 2, west), E (POP 3, east), C (POP 4, central) and S (POP 5, south).

Figure 4b and 4d show the dendrograms of male and female populations based on genetic (Cavalli Sforza and Edwards) distance. Two genetic groups were formed in the dendrogram of the male populations while no genetic groups were found in the female populations. The first group in the male genetic dendrogram consisted of mosquito populations from the west (POP2), east (POP3) and south (POP5) while the second group consisted of north (POP 1) and central (POP 4) populations. Bayesian analysis in GENELAND showed that the most probable number of genetically differentiated clusters using all male individuals across mosquito populations was *K* = 3 while in females, it was *K* = 2 (Figure S1).

### Comparison and Correlation between Genetic and Phenotypic Markers

The results generated by both phenotypic and genetic markers were compared via (a) *Q*_ST_ and *F*_ST_ (b) CV and DAPC plots (c) dendrogram topology and (d) Bayesian clustering. The estimated *Q*_ST_ was higher than *F*_ST_ in both male (0.318>0.055, p < 0.001) and female (0.309>0.009, p < 0.005) populations. In females, the CV and DAPC plots showed overlapping patterns among the populations. In contrast, both the CV and DAPC showed that all male populations were separated from each other. No groups were identified in the phenotypic and genetic dendrograms of the female populations. Both genetic and phenotypic dendrograms revealed two groups with nearly identical tree topologies in the male populations. Correlation analysis (Figure S2) revealed a significant association between the genetic and phenetic distances in the male populations (*r* = 0.68, *p* = 0.02) but not in the female populations (*r* = 0.27, *p* = 0.46). Bayesian analysis in the male and female populations detected multiple genetic clusters (*K* = 3 and *K*=2, respectively) while no phenotypic clusters (*K* = 1) were found in both populations.

## Discussion

### Phenotypic and Genetic variations between Female and Male *Ae. aegypti*

Overall, our results revealed contrasting phenotypic and genetic patterns between male and female *Ae. aegypti* populations, indicating the male populations to be spatially structured than female populations. This conclusion is supported by the increased variation among males, wing shape, DAPC plots and distinct dendrogram groupings for males only. The low genetic differentiation of female *Ae. aegypti* populations in Metropolitan Manila suggests a continuous and active exchange or sharing of alleles, indicating high gene flow and/or a weak genetic drift among female populations. The results imply that female *Ae. aegypti* passively disperse throughout the region. It is possible that *Ae. aegypti* take advantage of human-aided transportation land routes such as roads to travel to distant areas (Brown et al., 2014; Fonzi et al., 2015; Carvajal et al., 2020). The presence of eggs, larvae and adult mosquitoes in commercial trucks^45,46^ could also facilitate the high gene flow observed in our results. The wide spread movement of female *Ae. aegypti* mosquitoes can result in genetically-similar populations. This is consistent with the results of our study in that the female mosquito populations were genetically- and phenotypically-similar. These findings correlate well with a previous study in Metropolitan Manila where mosquito populations from the north, west and south of the region were deemed genetically-similar, indicating that road networks can facilitate passive dispersal of the mosquito vector^6^.

In contrast, the findings from male *Ae. aegypti* mosquitoes suggest limited gene flow and/or strong genetic drift. The limited gene flow could be due to the behavior and life expectancy of the male mosquito. We assume that males are unlikely to take advantage of human-mediated transportation to travel to distant locations. Males are considered to be less anthrophilic than their female counterparts since they do not need blood for reproduction or survival. Furthermore, males only swarm humans to mate with blood-seeking females, but mating was frequently observed at oviposition sites or breeding containers^47^. This specific behavior could restrict male *Ae. aegypti* mosquitoes in their movement to distant locations, thus encouraging the formation of heterogeneous population structures in the region.

Alternatively, the limited gene flow and/or strong genetic drift in males could be due to their short life span. The short life span of certain species can influence the connectivity (i.e. dispersal) of their populations, thereby affecting the distribution of population genetic variation and structure^48^. In addition, adult life-spans are directly related to effective population size, which can shape the genetic diversity of a population^49^. This is exemplified in short-lived populations of turquoise killifish^50^, where the resulting small effective population size was due to their short life span which led to a bottleneck effect in selected populations. In *Ae. aegypti*, females have a life expectancy of 10 to 35 days^51^ while that of males is of 3 to 6 days^52^. We thus infer that the shorter life span of male *Ae. aegypti* mosquitoes could result in a lower population size for a period of time, thereby strengthening genetic drift and producing distinct genetic structures among the male populations.

It should be noted that both markers (genetic and phenotypic) are heritable in nature. The female’s genetic and phenotypic characteristics (low spatial structure) can be inherited by the male offspring and *vice versa*. Therefore, the heritable nature of the markers may work to reduce the contradicting patterns between males and females over multiple generations^53^. Nevertheless, we observed clear sexual differences, suggesting that the population structures strongly reflected short-term population history occurring in a generation at the time sampling, rather than long-term or multi-generational history. We did not conduct any temporal investigation of these mosquito populations to confirm the steadiness or stability of the population structure in male mosquito populations. This avenue may be worth investigating in future endeavors.

### Comparing Phenotypic and Genotypic Approaches

Previous studies that used an integrative approach to study the spatial population structure of *Ae. aegypti* demonstrated contradictory findings between genetic and phenotypic markers^3,4,16,17^. In this study, opposing outcomes were observed such as (a) higher *Q*_ST_ than *F*_ST_ estimates and (b) higher numbers of detected clusters in genetic markers than phenotypic markers. Other analyses demonstrated near-mutually congruent results such as (c) similar contradicting patterns of males and females in the CV and DAPC plots and (d) similar dendrogram topologies and correlations between the estimated genetic and phenotypic distances.

Our results are consistent with past studies emphasizing that genetic markers (e.g., microsatellites) offer more sensitivity in detecting population variability and structure than wing shape^10,11^. If we compare the DAPC (genetic) and CV (phenotypic) plots, mosquito populations in DAPC plots are separated more than those in the CV plots. Nonetheless, even if the two types of plots were not identical in nature, both approaches revealed a similar interpretation where a more highly diverged population structure was found in the male populations, as compared to the female populations. It is also noteworthy that the near-identical groupings found on the dendrogram topologies were also found in the significant correlation analysis of the genetic and phenetic distances, especially in male mosquito populations. Although past studies yielded no significant correlation between the two markers, this particular finding was only found in female *Ae. aegypti* mosquitoes^3,4,16,17^. The likely reason for the correlating patterns between the two markers is that the genetic change at the level of point mutations and phenotypic mutations at the protein level were occurring concurrently at different mutation rates. Though phenotypic markers are considered to undergo evolutionary changes more slowly than genetic markers, there could be a certain level of phenotypic variation that was sustained and detected at the fine-scale due to neutral processes which could be occurring in parallel to both markers^54^. Thus, this study was able to detect these correlating patterns in the clustering and ordination analyses based on the population or individual distance. In addition, this correlating pattern between the two markers could also be due to the high spatial heterogeneity found in the genetic and phenotypic structures, which resulted from either isolation by distance or adaptive divergence^55,56^.

Previous studies have concluded that the absolute levels of variations (i.e., *F*_ST_ and *Q*_ST_) from these two approaches vary considerably for these two approaches. This is evidenced by the results of this study where *Q*_ST_ is larger than *F*_ST_ in both mosquito populations and a higher number of detected clusters were found in genetic markers (K= 2 – 3) than in phenotypic markers. This clearly indicates that phenotypic variability is low, suggesting that wing shapes among populations are homogenous or at least somewhat uniform^3,4,11,20^. Henry et al.^21^ theorized that two factors could account for the relative homogenous wing shape in *Ae. aegypti* namely; canalizing mechanisms and persistent gene flow. Canalization is a concept that illustrates the reduced sensitivity of phenotypes to genetic and environmental perturbance. Since wings are important for flight and mating behavior^3,57^, robustness in wing shape is expected at work. This is corroborated by our Bayesian analysis results, where multiple clusters were detected in the genetic analysis of both mosquito populations but no clusters were found in the phenotypic analysis. In addition, high gene flow due to continuous migrations of the mosquito vector can counteract local phenotypic drift processes. As mentioned previously, genetic markers demonstrated higher sensitivity in detecting population structures than the phenotypic markers did. This does not diminish the importance of using wing shape as a marker, as this type of marker can still serve as a preliminary tool in describing population structure and can be complemented by genetic markers that provide the necessary quantitative parameters to explain the evolutionary pattern of *Ae. aegypti*^11^.

## Conclusions

This study demonstrated the contrasting population variability and population structure between male and female *Ae. aegypti* populations using wing geometry and genetic analyses. Overall, male populations were more spatially structured than their female counterparts. *F*_ST_ estimates were higher in males (0.05) than in female (0.009) populations. Wing shape variation (CV plots) indicated that male populations are more separated. This pattern was also similar in the DAPC plots using genetic data. Genetic and phenetic dendrograms showed the formation of two groups of male populations but no groups were found in the female populations. In addition, Bayesian analysis using genetic data detected more genetic clusters in males populations (K= 3) than in female populations (K= 2).

## Additional Files

**Additional File 1: Table S1.** Sampling information for *Ae. aegypti* in Metropolitan Manila, Philippines

**Additional File 1: Table S2.** Comparative wing centroid sizes of each population in male and female *Ae. aegypti* mosquitoes

**Additional File 1: Table S3.** Morphological Metric Disparity (lower diagonal) and Mahalanobis distance (upper diagonal) across male and female *Ae. aegypti* populations

**Additional File 1: Table S4.** Allele frequencies of the 11 microsatellite loci in male and female *Ae. aegypti* populations in Metropolitan Manila

**Additional File 1: Table S5.** Analysis of the genetic diversity of male and female *Ae. aegypti* using 11 microsatellite loci

**Additional File 1: Table S6.** Null allele frequency estimates per locus of male and female *Ae. aegypti* populations

**Additional File 1: Table S7.** Analysis of linkage disequilibrium of male and female *Ae. aegypti* populations in Metropolitan Manila, Philippines

**Additional File 1: Table S8.** Genetic diversity of male and female *Ae. aegypti* populations based on 11 microsatellites in Metropolitan Manila, Philippines

**Additional File 1: Table S9.** Genetic pairwise *F*_ST_ (lower diagonal) and Cavalli-Sforza and Edwards distance (upper diagonal) across male and female *Ae. aegypti* populations

**Additional File 1: Table S10.** Analysis of molecular variance (AMOVA) of male and female *Ae. aegypti* populations from Metropolitan Manila

**Additional File 2: Figure S1.** Phenotypic and genetic clusters of male and female *Ae. aegypti* populations inferred with the program GENELAND. A bar plot of the density of each number of clusters along the chain after burn in. Patterns of population structure of *Ae. aegypti* inferred by the integration of georeferenced genetic/phenotypic data. Different colors indicate heterogeneous clusters.

**Additional File 3: Figure S2.** Scatterplot diagram of the phenotypic (Mahalanobis) and genetic (Cavalli Sforza and Edwards) distances in (a) female and (b) male mosquito populations.

## Acknowledgements

We would like to thank K.M. Viacrusis, L.F.T. Hernandez, H.T. Ho, M.J.L.B. Martinex, J.D.R. Capistrano and V.S.P. Tiopianco in the collection of adult mosquitoes. B.M.C. Orantia, C.R. Estrada and M.G. Cuenca in conducting preliminary geometric morphometric analysis. K. Ogishi and S. Yaegeshi in conducting preliminary genetic analysis.

## Declarations

### Ethical approval and consent to participate

Not applicable

### Consent for publication

Not applicable

### Consent for publication

All genetic data generated are deposited and available at vectorbase.org with the population biology project ID: VBP0000554. Phenotypic data (wing shape) is available upon request.

### Author’s Contribution

TMC, DMA and KW conceptualized and designed the study. TMC and DMA collected and identified the adult mosquito samples for the study. TMC conducted geometric morphometric and genetic analyses. TMC and KW wrote the manuscript.

